# Computational modelling unveils how epiblast remodelling and positioning rely on trophectoderm morphogenesis during mouse implantation

**DOI:** 10.1101/2020.06.08.140269

**Authors:** Joel Dokmegang, Moi Hoon Yap, Liangxiu Han, Matteo Cavaliere, René Doursat

## Abstract

Understanding the processes by which the mammalian embryo implants in the maternal uterus is a long-standing challenge in embryology. New insights into this morphogenetic event could be of great importance in helping, for example, to reduce human infertility. During implantation the blastocyst, composed of epiblast and trophectoderm, undergoes significant remodelling from an oval ball to an egg cylinder. A main feature of this transformation is symmetry breaking and reshaping of the epiblast into a “cup”. Based on previous studies, we hypothesise that this event is the result of mechanical constraints originating from the trophectoderm, which is also significantly transformed during this process. In order to investigate this hypothesis we propose MG#, an original computational model of biomechanics able to reproduce key cell shape changes and tissue level behaviours *in silico*. With this model, we simulate epiblast and trophectoderm morphogenesis during implantation. First, our results uphold experimental findings that repulsion at the apical surface of the epiblast is sufficient to drive lumenogenesis. Then, we provide new theoretical evidence that trophectoderm morphogenesis indeed dictates the cup shape of the epiblast and fosters its movement towards the uterine tissue. Together, these results offer mechanical insights into mouse implantation and highlight the usefulness of agent-based modelling methods in the study of embryogenesis.

**Author summary:** Computational modelling is increasingly used in the context of biological development. Here we propose a novel agent-based model of biological cell and tissue mechanics to investigate important morphological changes during mouse embryo implantation. Our model is able to replicate key biological cell shape changes and tissue-level behaviour. Simulating mouse implantation with this model, we bring theoretical support to previous experimental observations that lumenogenesis in the epiblast is driven by repulsion, and provide theoretical evidence that changes in epiblast shape during implantation are regulated by trophectoderm development.

## Introduction

A critical milestone of mammalian development is reached when the embryo implants in the maternal uterine tissue [1, 2]. Prior to implantation, a series of cell fate decisions concomitant with multiple rounds of divisions gradually transform the initial zygote into a blastocyst featuring three different cell lineages: a spherical embryonic epiblast (EPI) wrapped into two extraembryonic tissues, the trophectoderm (TE) and primitive or visceral endoderm (PE/VE) [3, 4]. Upon implantation, the embryo moves towards maternal sites, and undergoes significant remodelling, culminating in the case of the mouse in an egg cylinder, a body structure essential to post-implantation phases such as gastrulation [4–6]. A key feature of this blastocyst-to-egg-cylinder transition, still poorly understood, is the appearance of symmetry breaking within the epiblast and its reshaping into a cup [4, 7], which occurs roughly between stages E4.5 and E4.75 of embryonic development.

Many of the important structural changes that occur during implantation have been explained in terms of chemical signals within and between embryonic and extraembryonic compartments [1, 8]. For instance, it was shown that at the onset of implantation epiblast cells exit their naive pluripotency state, self-organise into a highly polarised rosette, and initiate lumenogenesis under the influence of *β*1-integrin signalling [7, 9]. Shortly after implantation, *β*1-integrin enables pro-amniotic cavity formation along the entire egg cylinder via the resolution of multiple rosettes both in extraembryonic cell populations and at their interface with the embryonic tissue [6]. Moreover, differentiation of the primitive trophectoderm into polar and mural trophectoderm leading to the formation of a boundary between the two tissues was traced back to fibroblast growth factors (FGFs) signalling [10].

As D’Arcy Thompson already noted about genetics, however, development cannot be construed solely in terms of biochemical signals either: the mechanical interactions between cells and tissues equally and reciprocally contribute to embryogenesis [11, 12]. On the subject of the epiblast remodelling into a cup, a series of biological works have paved the way and triggered further investigation into the mechanics involved. Because it was observed that the EPI did not initiate specific tissue-level symmetry-breaking behaviours, one study stated that after the basement membrane disintegrated between the EPI and TE, the membrane between the EPI and the PE acted like a basket that moulded the epiblast into its cup shape [4] (Fig. 1A). Although this hypothesis put the spotlight on the basement membrane, it also suggested that the TE in direct contact with the EPI could play a role in this shape change. Evidence supporting this hypothesis grew when “ETS-embryoids” (ETS: embryonic and trophoblast stem-cell) assembled *in vitro* from EPI and TE stem cells, surrounded by the extracellular matrix (ECM) acting as the basement membrane, replicated embryonic transition from blastocyst to egg cylinder [13] (Fig. 1B). Furthermore, a recent study highlighted more clearly the role of the trophectoderm [14]. In this study, ExE-embryoids (ExE: extra-embryonic ectoderm), cultured from EPI and PE stem cells separated by an ECM basement membrane, did not break the symmetry of their initial spherical shape (Fig. 1C). In contrast, both ETS- and ETX-embryoids (ETX: embryonic, trophoblast and extra-embryonic endoderm) made from all three blastocyst lineages did reproduce the symmetry breaking observed in real embryos. Together, these studies established the necessity of the trophectoderm for the remodelling of the epiblast [13, 14].

**Fig 1.**
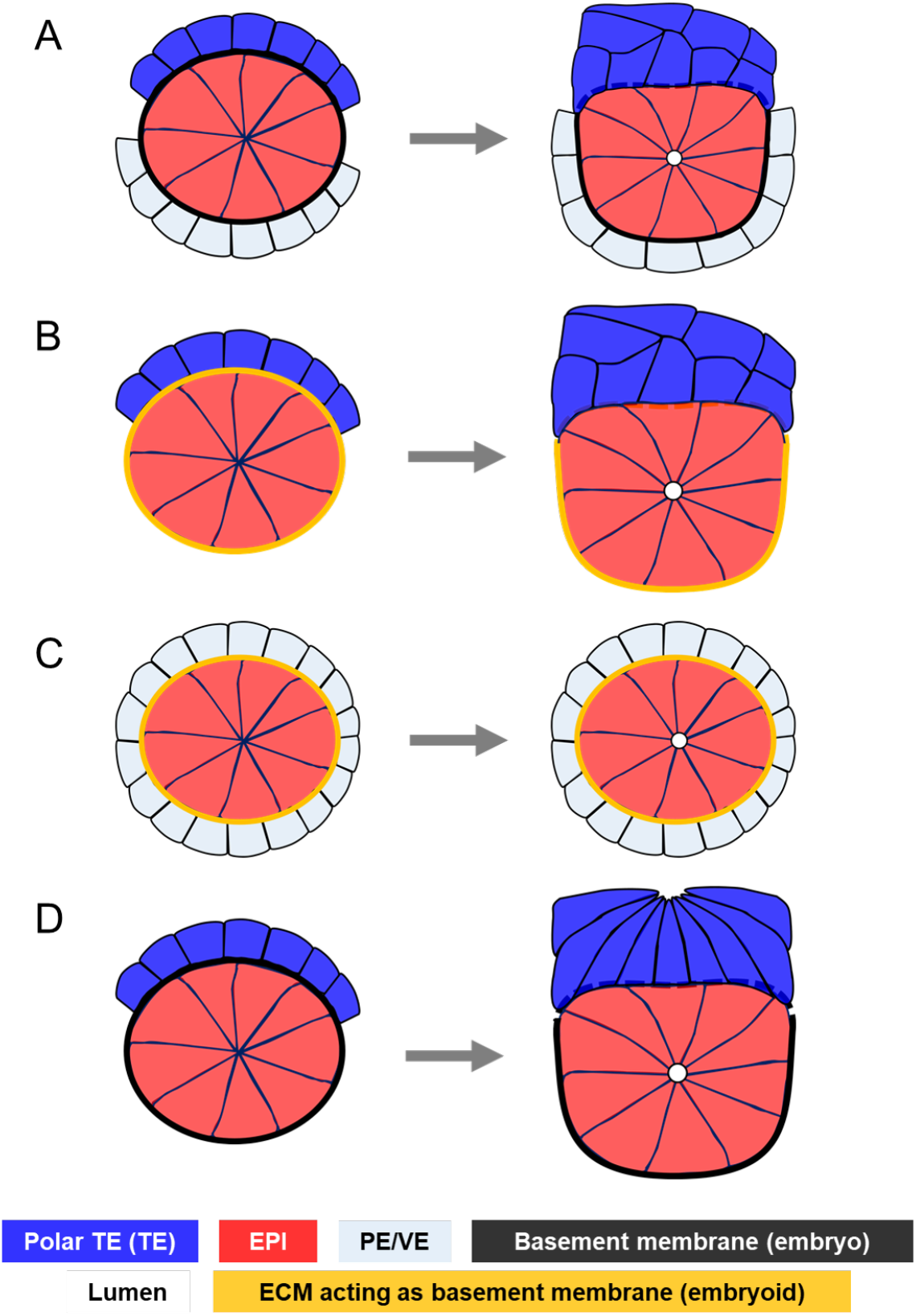
Review of epiblast symmetry breaking theories. **A.** The basement membrane separating the epiblast and the primitive endoderm moulds the epiblast into a cup while it disintegrates between the epiblast and the trophectoderm in mouse embryos [4]. **B.** Embryoid structures featuring epiblast and trophectoderm stem cells surrounded by an ECM acting as a basement membrane (ETS-embryoids) replicate mouse embryogenesis by forming body structures similar to those observed in normal embryonic development [13]. Here the presence of the trophecdoderm shows that this tissue might be required for symmetry breaking in the epiblast and cup shape acquisition. **C.** Embryoid structures featuring epiblast and primitive endoderm stem cells surrounded by an ECM acting as a basement membrane (EXE-embryoids) do not break symmetry in the epiblast, but initiate lumenogenesis [14]. This evidences the requirement of the trophectoderm for the remodelling of the epiblast. **D.** Trophectoderm morphogenesis during mouse implantation. Trophectodermal cells elongate, then undergo apical constriction, resulting in the tissue folding and invaginating the epiblast [10]. This suggests that epiblast remodelling into a cup might be a mechanical response to trophectoderm dynamics

On the other hand, *how* exactly trophectoderm morphogenesis influences shape change in the epiblast has not been elucidated yet because very little is known on trophectoderm morphogenesis during implantation. In the light of recent detailed descriptions of extra-embryonic tissues morphogenesis during implantation [10], it appears increasingly plausible that trophectoderm morphogenesis regulated epiblast remodelling via mechanical interactions at their common boundary. This study showed that polar trophectodermal cells exhibited drastic morphological changes throughout the implantation period. Whereas early implanting blastocysts featured squamous cells in the polar trophectoderm, these cells, driven by a high mitotic and space restrictions due to the formation of a boundary with the mural trophectoderm, later transited to cuboidal, then elongated to acquire columnar shapes. These changes were followed by apical constriction resulting in the folding of the whole tissue, and invagination of the epiblast (Fig. 1D). Moreover, this study provided experimental evidence that other structural changes, most notably the stretching of PE cells, resulted from TE morphogenesis [10]. Hence, we want to investigate the hypothesis that trophectoderm morphogenesis drives the remodelling of the epiblast into a cup via mechanical interactions at their common boundary.

Building on the increasing power of computational modelling in developmental biology [15–18], we examine the influence of trophectoderm morphogenesis on the epiblast. The requirement of dramatic cell shape changes in trophectodermal cells, notably apical constriction [10], orients modelling options toward the family of deformable cell models (DCM) [19]. In this category, two classes of models have been predominant in recent research: vertex models (VM) and sub-cellular element models (SEM). Although vertex models were used extensively to study epithelial dynamics [20, 21], accounting for various mechanical behaviours of individual cells remains challenging in a global energy-based approach. Hence, we set our choice on SEM, where cells are represented by an agglomeration of computational particles interacting with one another via short-range potentials emulating the viscoelastic properties of their cytoskeleton [22–24]. However, in order to exhibit realistic cell shapes, SEM generally involve an important number of particles, many of which reside within the cell, thus do not have a direct influence on cell shape. This leads to increased computational complexity, limiting the size of cell populations that can be simulated.

Here, we present a novel computational SEM called MG#, which focuses on 3D cell shapes while reducing computational complexity by distinguishing between membrane particles and a single intracellular particle. Using this model, we first uphold the experimental observation that repulsion at the apical surface is sufficient for lumenogenesis in the epiblast. Then, we reproduce trophectoderm morphogenesis during implantation and we provide theoretical support that epiblast remodelling into a cup shape and its movement towards the maternal uterine tissue can be explained by trophectoderm morphogenesis.

## Model

Based on the fundamental principles of DCM, our abstraction of the biological cell features particles in interaction under the influence of potential-derived forces. Emphasis is put on particles at the surface of the cell membrane, bringing our model close to VM [25], while at the same time we also include a single intracellular particle reminiscent of the cell’s microtubule organising centre (Fig. 2A,B).

**Fig 2.**
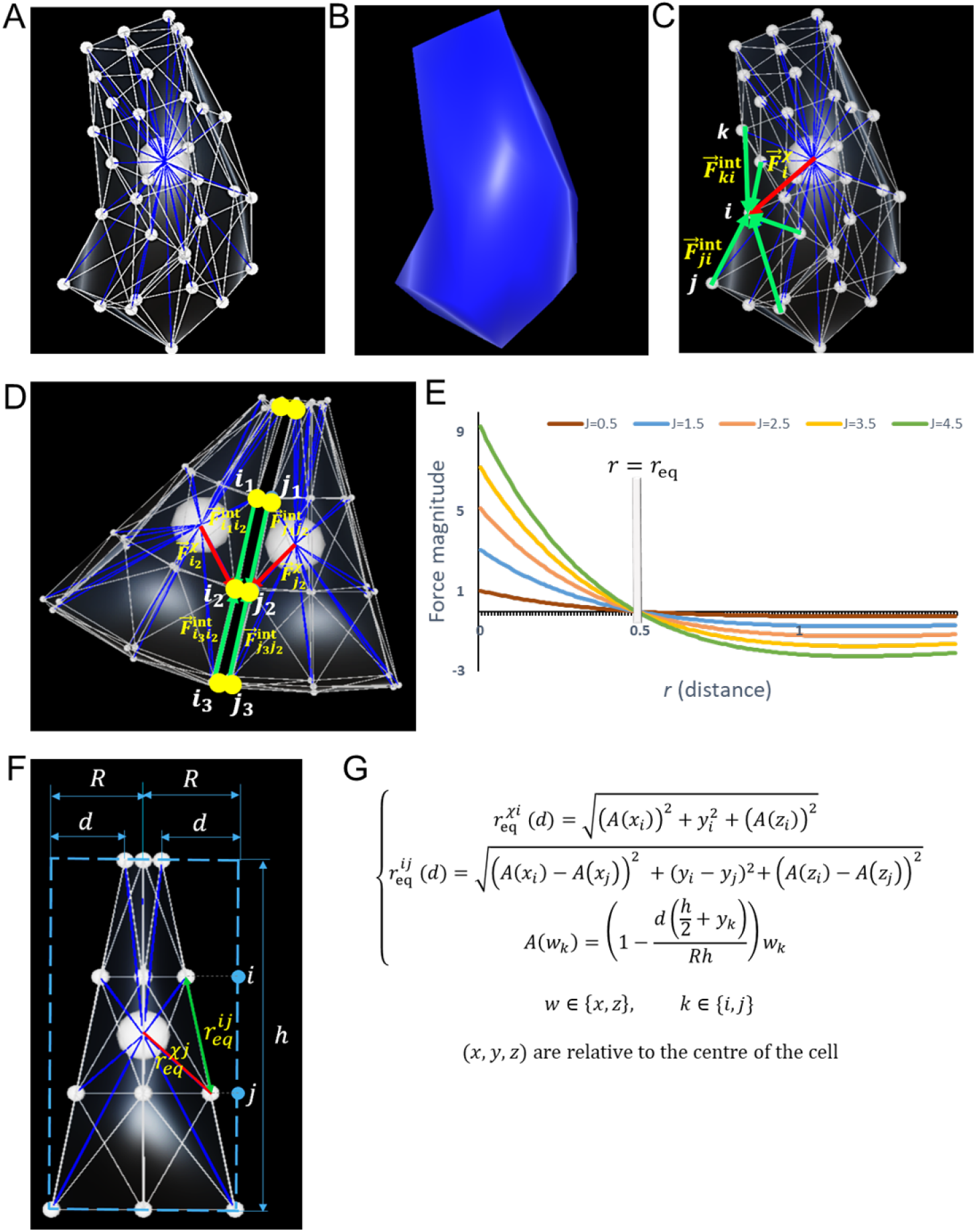
Computational model. **A.** 3D representation of a cell: The cell is abstracted by an agglomeration of particles (small white spheres), whose triangulation (white edges) forms the membrane, and by an intracellular particle (big white sphere). Interactions between the intracellular and membrane particles (blue lines) mimic the cytoskeleton. **B.** 3D rendering of a cell without its sub-cellular elements. **C.** Forces acting within a cell: 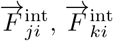 are the forces that membrane particles *j, k* exert on another membrane particle *i*. 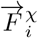 is the force that the intracellular particle *χ* exerts on *i*. **D.** External forces acting on a cell via its particles. Here, 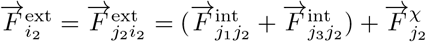. **E.** Plots of the magnitude of Morse forces under different values of *J*, with *ρ* = 1 and *r*_eq_ = 0.5. **F.** Apical constriction of an epithelial cell with original radius *R* shrinking by *d*. **G.** Formulas of the new equilibrium lengths in an apically constricted cell.

On the cell membrane, we define a topological neighbourhood based on a triangulation of vertices. Two same cell particles are deemed internal neighbours if they both belong to one of the mesh triangles (Fig. 2A). We also define an external neighbourhood based on distances between particles of different cells (Fig. 2D). To minimise the computation time required, we introduce cell-cell neighbourhood relationships where particles of different cells are tested for external neighbour links only when the cells to which they belong were already approved as neighbours. Here, a Moore neighbourhood, well suited for the lattice-like layout of our cells, is favoured.

In order to induce intrinsic mechanical behaviours within cells, we assimilate internal particle neighbourhood links to non-linear springs, which have been shown to faithfully emulate living matter [26]. These springs mimic the activity of actomyosin and microtubule networks in the cytoskeleton, and forces are derived from their elastic potential (Fig. 2C-E). In the cell’s resting state, the equilibrium distance of each spring coincides with the length of the segment formed by its nodes. Cell dynamics arise from alterations to these equilibrium distances. In apical constriction for instance, new equilibrium lengths are computed as in Fig. 2F,G.

### Equation of motion

Acting on a given membrane particle *i*, we distinguish four main types of forces: internal forces 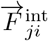, cytoskeleton forces 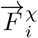, external forces 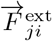, and specific forces 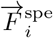. Biological media are often characterised by a low Reynolds number, due to their high viscosity, which minimises the effects of inertia [19]. We therefore subject particles to an over-damped, first-order equation of motion:

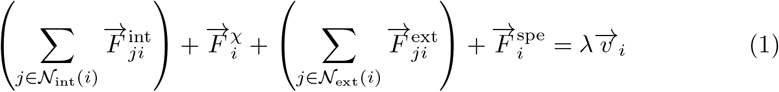

where 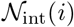 and 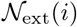 represent the sets of internal and external neighbours of particle *i*, and λ is the coefficient of friction exerted on all particles.

### Internal and cytoskeleton forces

The internal force created by a particle *j* on a neighbouring particle *i* derives from a Morse potential (Fig. 2E). Previous studies have used Morse potentials to represent forces in a biological context [22, 24]. The expression of this force is given by:

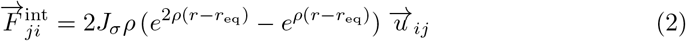

where *J_σ_* represents the interaction strength between particles *i* and *j*, both of cell type *σ*, *r*_eq_ is the equilibrium of the spring force between *i* and *j*, and 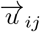 is the unit vector along the direction formed by *i* and *j*. Similar forces dictate interactions between the intracellular particle and the membrane particles.

### External forces

Given the tight packing in epithelial tissues, a cell membrane is always in contact with neighbouring cell membranes. Thus local action on a membrane produces an equivalent deformation on the surrounding cells. In other words, a particle always transmits the force received to its external neighbours. To account for this behaviour, we submit particles and their external neighbours to equal forces. This is done by setting the external force acting on a particle to be equal to the sum over all its external neighbours of their internal and nucleus forces:

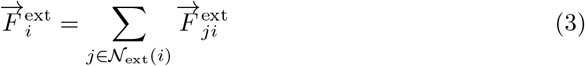

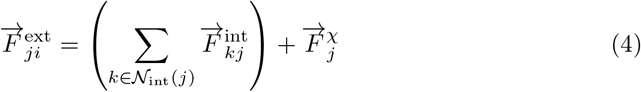

### Specific forces

Generally speaking, it is possible to include specific forces in DCM to account for desired behaviours. A few studies have taken advantage of this possibility to enable for example cell surface bending resistance [26] or cell surface area and volume conservation [27]. In our context of mouse implantation morphogenesis, we introduce specific forces to simulate repulsion at the apical surface of epiblast cells during lumenogenesis [4, 9, 14]. Here, these forces also derive from a Morse potential.

## Results

In this section, we applied our model to the study of mouse embryo morphogenesis during implantation. Here we focused on epiblast and trophectoderm tissues. First, we tested the hypothesis of whether repulsion at the apical surface of the epiblast was sufficient to account for lumenogenesis. Then, we simulated both tissues’ morphogenesis and showed that the epiblast remodelling into a cup shape and its movement towards the maternal uterine tissue could be explained by trophectoderm morphogenesis.

Simulations were run using a C# implementation of the model described above. The source code for the simulation engine can be found at https://github.com/guijoe/MGSharpCore. The repository for the Unity3D-based viewer can be found at https://github.com/guijoe/MGSharpViewer.

### Repulsion at the apical surface of the epiblast is sufficient for lumenogenesis

The study of how lumens arise in epithelial tissues has revealed two predominant mechanisms: cavitation mediated by apoptosis, and hollowing, in which the lumen is formed by exocytosis and membrane separation [28, 29]. In the case of highly polarised epithelia, it was shown that cavitation was not necessary for lumenogenesis [30]. Hence, the hollowing mechanism was privileged in epiblast lumenogenesis, which features highly polarised cells spatially organised in the shape of a rosette. It was hypothesised that charge repulsion mediated by anti-adhesive molecules such as podocalyxin (PCX) drove lumen formation in the epiblast [4, 7]. Furthermore, evidence for hollowing was observed in a recent study [14], where apoptosis was found not to regulate lumenogenesis, but PCX was discovered to be predominant at the apical surface of cells facing the lumen.

Using our model, we sought to determine theoretically whether hollowing via repulsion at the apical surface of the epiblast rosette was a viable mechanism for lumenogenesis in this tissue. First, we built a 3D rosette-shaped epiblast by submitting polarised epithelial cells to apical constriction [7] (Fig. 3A,B, Supplementary Fig. S1A). Then, inspired by the anti-adhesive role of PCX, we broke adhesive links between appropriate cell membranes in contact at the apical surface of the rosette, and created repulsive forces (Eq. 2). This prompted neighbouring apical particles to break apart from each other, initiate and gradually expand a lumen at the centre of the rosette (Fig. 3C-E). This result therefore suggests that hollowing via repulsion is sufficient as a mechanism for lumenogenesis in the mouse epiblast.

**Fig 3.**
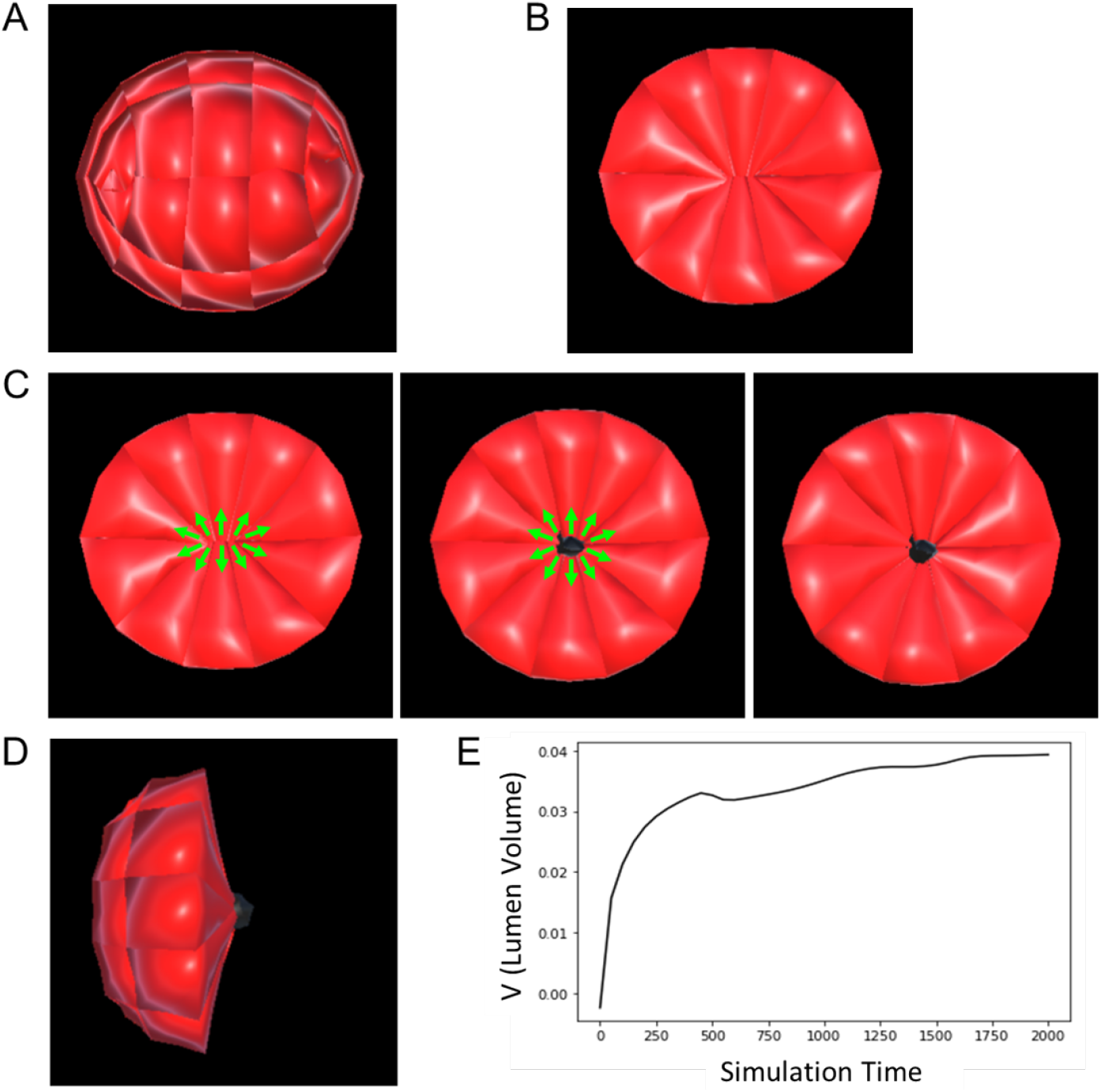
Lumenogenesis in the epiblast. **A.** A 3D model of a rosette-shaped epiblast. **B.** A 2D slice of the epiblast in **A** showing apically constricted cells of the building block of the epiblast rosette. **C.** Creation of the lumen cavity by repulsion at the apical surface of the epiblast. Green arrows represent the direction of repulsive forces. The snapshots (from left to right) were taken respectively at *t* = 0, 500 and 2000. **D.** Lateral view of the sliced epiblast showing the lumen volume. **E.** Dynamics in time of the volume of the lumen. Values of the equation parameters: *J*_EPI_ = 2:5, λ = 2, *ρ* = 1.

### Mechanical constraints imposed by TE morphogenesis on the epiblast drive cup shape acquisition

A key feature of the blastocyst-to-egg-cylinder transition is the symmetry breaking within the epiblast and its shaping into a cup [4, 7]. During this transformation, the epiblast remodels from an oval ball to a tissue with a flat surface at its boundary with the trophectoderm. Previous studies have established the requirement of the trophectoderm in this shape change [13, 14]. Using the presented model, we investigated how trophectoderm morphogenesis influenced the cup shape acquisition by the epiblast. Our simulation protocol consisted of reproducing the sequence of morphological events observed in the trophectoderm as described in [10] (elongation followed folding via apical constriction), and keeping track of the consequent changes in the epiblast. For simplicity and to keep the model computationally efficient, we assumed that there were no cell divisions in the tissue.

We built a virtual embryo consisting of a TE sheet with initial cuboidal cells laying on top of an oval rosette-shaped epiblast (Supplementary Fig. S1B). At the initial stage (Fig. 4A,E), new equilibrium lengths were computed for all TE cells, with the goal of triggering a transition from cuboidal cells to more elongated columnar shapes. These cells lost their resting state and regained it by gradually aligning their actual springs lengths with the calculated equilibrium lengths (Fig. 4B,F). After that, we initiated invagination in the TE. The distribution over the entire sheet of the length *d* by which the apical radius of cells *r* was shrunk depended on the position of the cell in relation to the centre of the sheet. In our simulations, this distribution was given by a step function: cells in the middle of the sheet were set to constrict completely (*d* = *r*), while cells on the boundary did not constrict (Supplementary Fig. S2). The coordinated movement of cells induced by these positional laws caused the tissue to fold and invaginate the epiblast. Short after TE invagination begins, we initiated lumenogenesis in the epiblast (Fig. 4G). In order to highlight the requirement of the TE, following TE folding (Fig. 4C,G), we broke the contacts between the TE and the epiblast for the remaining time of the simulation, inhibiting any mechanical interactions between the two tissues, but maintaining both tissues’ own mechanics (Fig. 4D,H). We noted that throughout the experiment, with the exception of lumenogenesis, epiblast cells did not initiate any behaviours, the epiblast as a whole simply reacted to the mechanics induced by either the presence or the absence of the TE.

**Fig 4.**
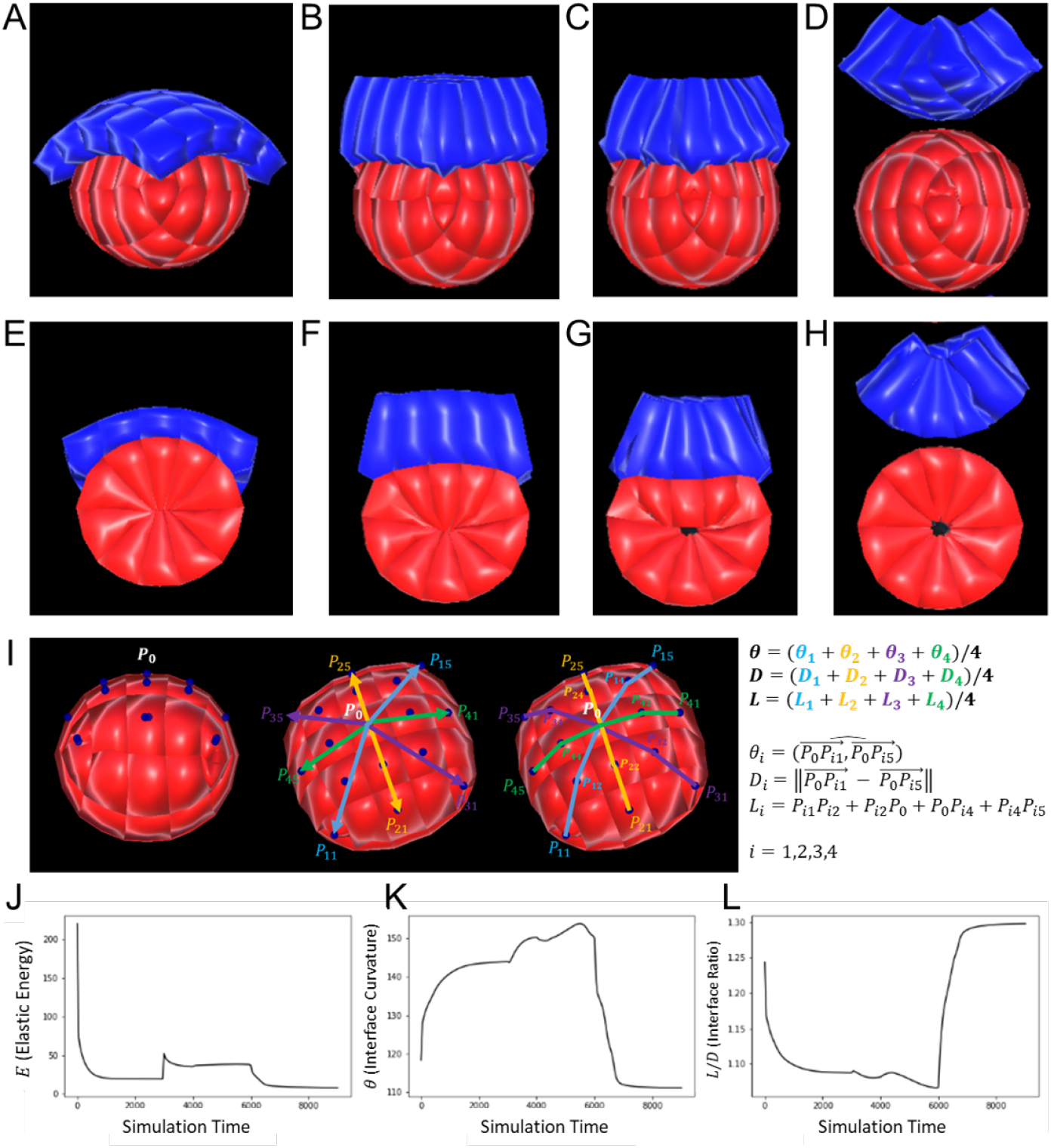
Trophectoderm morphogenesis regulates epiblast shape. **A-D.** 3D snapshots of the simulation of TE and EPI morphogenesis during mouse implantation, and the regulation of EPI shape, taken respectively at *t* = 0, 3000, 6000 and 9000. **E-H.** Corresponding 2D slices of the cell population at the same stages. **(A,E).** The initial stage features a single layered TE with cuboidal cells resting upon the rosette-shaped epiblast. **(B,F).** TE cells have transited to a columnar shape. **(C,G).** The TE has folded by apical constriction of single cells. Concomitantly, lumenogenesis was initiated in the epiblast (the process starts at *t* = 4000). **(D,H).** After adhesive links were broken between TE and EPI, the EPI bounces back to its near spherical shape. **I.** Definitions of the metrics used to evaluate the model, involving the curvature *θ*, TE/EPI interface diameter *D*, TE/EPI interface length *L*, and interface ratio *L/D*. **J.** Plot of the population’s elastic energy *E*. Discontinuities mark the start of new morphological events at *t* = 0, 3000, 4000, and 6000). After removal of the TE, *E* falls closer to zero than ever before, meaning that cells are closer to their resting stage, hence less externally constrained. **K.** Plot of the interface curvature *θ*. During TE morphogenesis, *θ* rises towards a flat angle, then sharply drops when the TE is removed. **L.** Plot of the interface ratio *L/D*. During TE morphogenesis, the interface curvature decreases towards 1, then sharply increases when the TE is removed. Values of the equation parameters: *J*_EPI_ = *J*_TE_ = 2.5, λ = 2, *ρ* = 1.

To appreciate the impact of the TE on the epiblast, we defined the elastic energy *E_i_* of a cell *i* as the sum over all cell springs of the squared difference between equilibrium and actual lengths. We extended this notion by defining the total elastic energy of a tissue or an entire population of cells as the sum of *E_i_*’s in the population. Cells always tended to minimise this energy, which can also be viewed as the degree of relaxation of cell: the closer it is to zero, the closer the cell is in its resting state, the more relaxed it is, hence the less constrained. In addition, we monitored the curvature of the epiblast, i.e. the inclination angle *θ* of the epiblast surface covered by the trophectoderm (Fig. 4I). An increasing curvature, trending towards a flat surface, was characteristic of the epiblast’s transition from an oval rosette to a cup. Moreover, we measured the length *L* and diameter *D* of the interface between EPI and TE, and considered their interface ratio *L/D* as our third evaluation metric (Fig. 4I). It was expected that this ratio would decrease towards 1 as the epiblast flattened. We plotted the profiles of the curvature, the interface ratio and the elastic energy throughout our simulation.

Our model matched biological expectations by replicating, on the one hand, an increasing curvature and a decreasing interface ratio, with ultimately a flat TE/EPI interface just before we removed the TE (Fig. 4C,G,K,L). On the other hand, as soon as the TE was removed, the epiblast bounced back to its original shape (Fig. 4D,H,K,L). This result agrees with the experimental observation that without the TE, the epiblast does break symmetry [14]. The elastic energy profiles tie these behaviours to the mechanical influence of the TE over the epiblast. Actually, breaking mechanical interactions between the TE and the EPI not only resulted in a sharp drop in elastic energy, but this energy also plateaued at a value significantly lower than in other stages (Fig. 4J), demonstrating that cells were more mechanically constrained when both tissues were in contact.

These observations suggest that the presence of the TE imposes mechanical stress on epiblast cells, hinting to the necessity of this tissue’s morphogenesis in the remodelling of the epiblast.

### Trophectoderm morphogenesis fosters epiblast movement towards the uterine tissue

An important requirement of implantation is close contact between the embryo and the uterine tissue. As soon as the three pre-implantation lineages are specified, the blastocyst hatches out of the zona pellucida and initiates the process of implantation [4]. However, there exists a gap between the hatched blastocyst and attachment sites in the uterus. In order to close this gap, the embryo needs to move towards the uterus. It was recently established that this movement of the embryo towards maternal sites occur concomitantly to the drastic morphological changes observed in the TE [10]. Furthermore, it was observed in that same study that primitive endoderm expansion over the whole embryo is driven by TE morphogenesis. Given that the trophectoderm keeps close contact with the epiblast during these events, we hypothesised that epiblast positioning could also be affected by TE morphogenesis. We employed computational modelling to examine whether TE morphological changes could influence the trajectory of the epiblast.

Here, as previously, we reproduced the sequence of TE morphogenesis (elongation followed folding via apical constriction), and observed how it affected the position of the epiblast. To highlight how the TE influences the trajectory of the epiblast, we defined what we designated as the “pushing distance”. We computed this distance at any given time point of the simulation by calculating the difference in height between the lowest point of the epiblast at that time point and the lowest point at the initial stage (Fig. 5A). We plotted the profiles of this metric and observed an increasing pushing distance as the TE transited from cuboidal to columnar, then as the TE folded (Fig. 5B). The sudden soar observed at *t* = 4000 reflects the slight elongation of the tissue due to hollowing-driven lumenogenesis in the epiblast.

**Fig 5.**
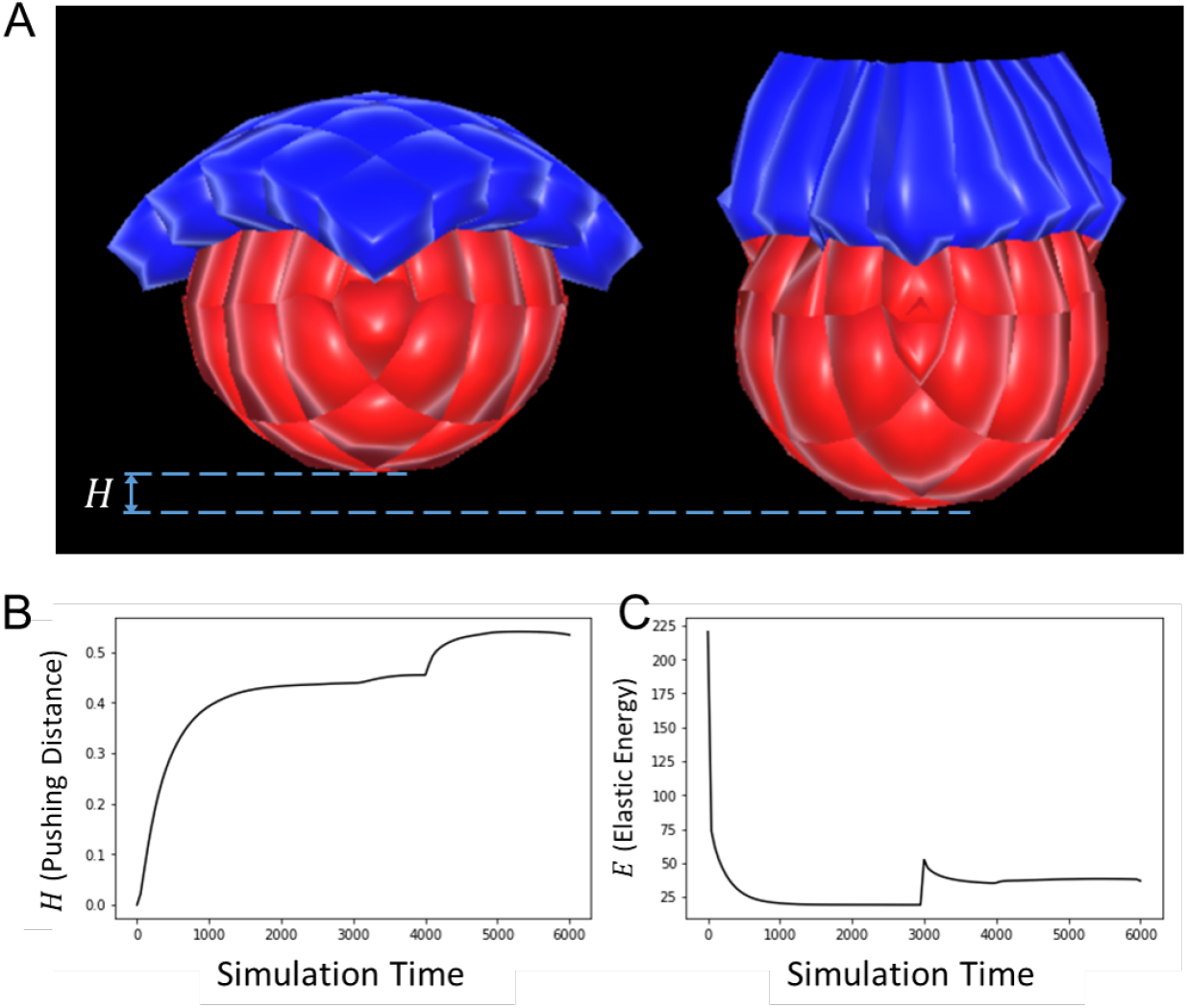
Trophectoderm fosters epiblast movement towards maternal sites. **A.** Snapshots of the simulation of TE and EPI morphogenesis during mouse implantation, and their influence on EPI positioning, taken respectively at *t* = 0 and 6000. **B.** Plot of the pushing distance, which increases with time. **C.** Plot of the elastic energy *E*. Discontinuities mark the start of new morphological events (*t* = 0 and 3000). The sudden soar observed at *t* = 4000 reflects the slight elongation of the tissue due to hollowing-driven lumenogenesis in the epiblast. Values of the equation parameters: *J*_EPI_ = *J*_TE_ = 2.5, λ = 2, *ρ* = 1.

These results suggest that TE morphogenesis, while reshaping the epiblast, also fosters the embryo’s movement towards maternal sites.

## Discussion and Conclusion

Understanding the processes by which the mammalian embryo implants in the maternal uterus is crucial to many breakthroughs in embryology [1]. New insights into these morphogenesis events could be of great importance in helping for example to reduce human infertility [31]. Although advances have been made by studying biochemical cues involved in these events, we focused here on the mechanical basis at the cellular level of epiblast morphogenesis. In order to study the physical dynamics of mouse implantation, we have designed a novel, computationally efficient model of biological cells and tissue mechanics able to simulate key episodes of vertebrate morphogenesis. With this model, we were able to schematically reproduce lumenogenesis in the epiblast, trophectoderm morphogenesis driven by single cells elongation and apical constriction, as well as provide theoretical support to the fact that this morphogenesis regulates the remodelling and positioning of the epiblast during implantation.

A well-known shortcoming of agent-based modelling is the risk to introduce disputable artefacts in the simulations. Within the scope of this work, we have shown that our model adhered well to biology by successfully simulating tissue-level morphological changes based solely on changes triggered at the cellular level, in a bottom-up, emergent fashion. We did this in particular for epithelial bending through apical constriction [32], rosette formation via polarised apical constriction [33], and repulsion-driven lumenogenesis [4, 7]. Nonetheless, some nuance should be added to certain quantitative features of the simulations. For instance, although it is a biological fact that the epiblast lumen’s volume increases as a result of cells drifting apart, the rate of this growth as exhibited in the graph of Fig. 3E may not reflect the actual rate curve in mouse embryos. The same could be said of the rate at which the epiblast reshapes (Fig. 4K,L), or the trophectoderm-induced epiblast velocity in its motion towards maternal sites (Fig. 5B). While not invalidating our main conclusions, these quantitative outputs are essentially contingent upon the choice of the potential function (here the Morse potential) and parameter values. This limitation could be overcome by experimenting with other potential functions, searching parameters space, and comparing results against real biological data.

Another weakness of computational modelling is its inability to integrate all possible details of a real-world problem, as this would inevitably increase complexity and demand unavailable computing power. Clearly, efficiency in our simulations was achieved by stripping the model of noticeable features of biological development. One important approximation is that we ignored the hypothetical impact of proliferation, although it is a pervasive phenomenon in both tissues. However, while it may be argued that it plays a non-trivial role in the elongation of trophectodermal cells [10], it is difficult to make a case that proliferation would be central in reshaping the epiblast. In fact, this particular lack in our approach could even be considered an advantage since neglecting proliferation also allowed isolating, hence highlighting the effects of pure mechanical interactions within and between the trophectoderm and the epiblast. Another simplification is that we neglected stochastic effects related for example to cell movements or the timing of embryogenesis episodes. To address this problem, stochastic models like Monte Carlo methods could be used to simulate single-cell apical constriction. Nevertheless, deterministic models, represented in this case by definite analytical equations, still exhibit good predictive power while remaining computationally practical.

In summary, although relatively abstract and schematic, our computational model and simulations offer new insights into mouse embryo implantation. Looking forward, refinements could combine the effects of mechanical interactions with proliferation and the stochasticity of biological processes to further investigate tissue shape changes. Then, the variables and parameters in these simulations could be tuned to fit quantitative metrics based on real measurements gathered from implanting embryos.

## Acknowledgments

JD acknowledges stimulating and fruitful discussions with Antonia Weberling, Ewa Paluch and Magdalena Zernicka-Goetz.

## Supporting information

**Fig S1.**
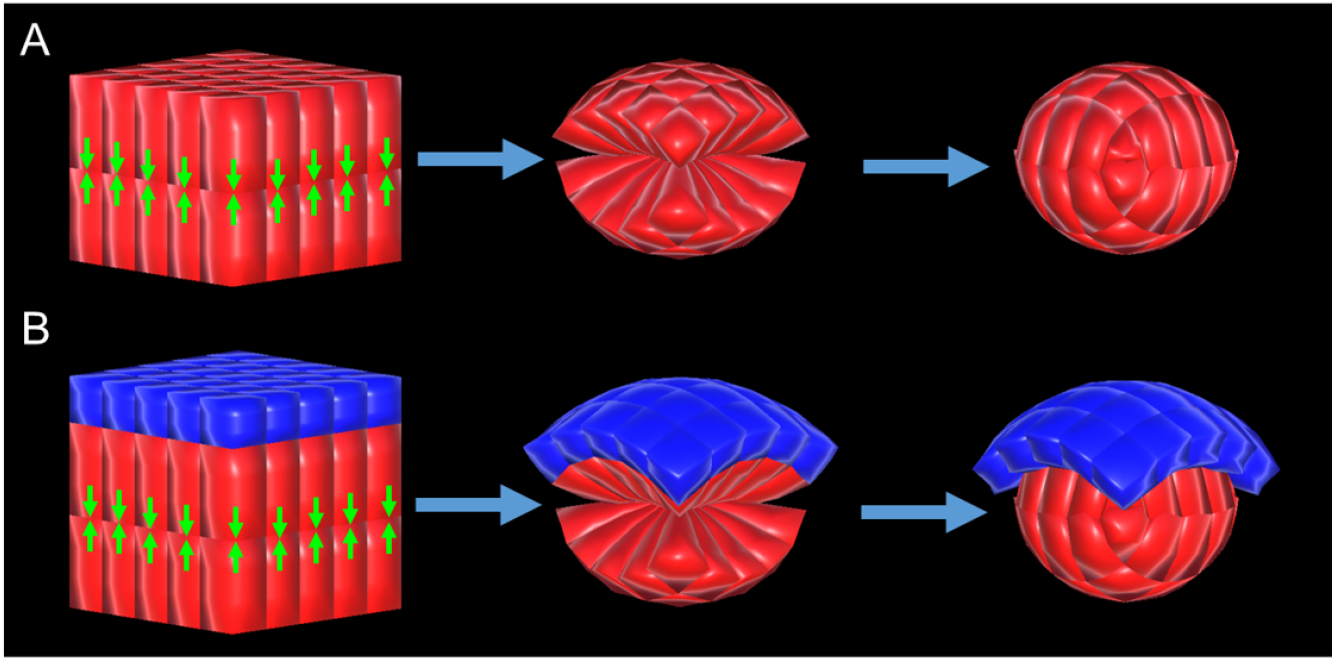
Epiblast and trophectoderm population reconstruction. **A.** The rosette-shaped EPI tissue is built by submitting polarised cells in a double epithelial layer to apical constriction. Green arrows indicate the apical surface of the cells, where constriction occurs. **B.** The initial cell population (TE and EPI) is built by adding an epithelial layer to the forming the EPI.

**Fig S2.**
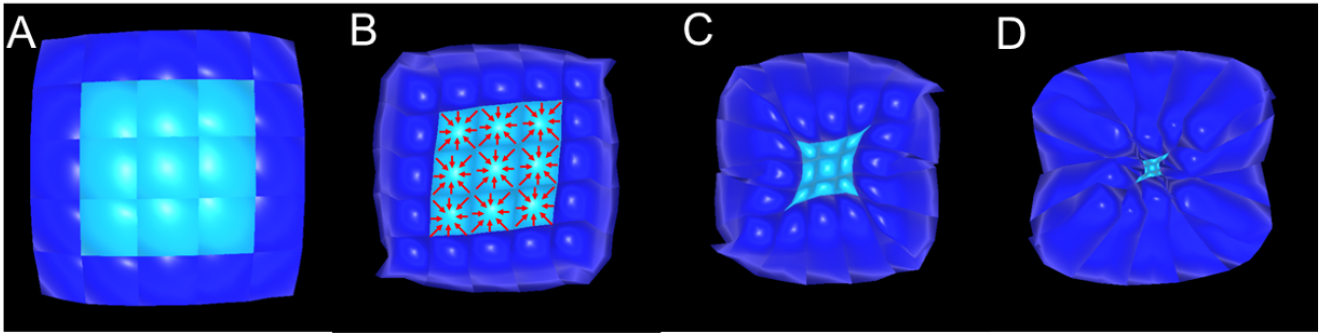
Top view of trophectoderm morphogenesis. **A.** Initial stage with cuboidal cells. **B.** Columnar TE initiating apical constriction. Red arrows highlight cells undergoing apical constriction. In this case, only cells in the middle constrict (light blue) to enable folding. **C.** Folded TE. **D.** Folded TE after separation from the EPI.

**Mov S3. Simulated morphogenesis during mouse implantation.** Trophectoderm cells elongate and then undergo apical constriction, leading the tissue to fold. At the same time, the epiblast remodels from a nearly spherical tissue to a cup-shaped tissue, while also undergoing lumenogenesis.

**Mov S4. Trophectoderm regulates epiblast shape.** Trophectoderm and epiblast undergo their normal development sequences (signle cells elongation followed by folding of trophectoderm, and reshaping and lumenogenesis in the epiblast). After the trophetoderm is detached from the epiblast, the epiblast bounces back to its nearly spherical shape. This shows that the epiblast broke symmetry and remodelled in the first place under mechanical stress imposed by trophectoderm morphogenesis.

